# Discriminative learning of substitution matrices and gap penalties for pairwise alignment of biological sequences

**DOI:** 10.64898/2026.05.14.725168

**Authors:** Michał Aleksander Ciach, Elissavet Zacharopoulou, Michał Piotr Startek, Błażej Miasojedow, Panagiotis Alexiou

## Abstract

Pairwise alignment scores are used to classify pairs of sequences in many areas of bioinformatics, including homology search, predicting interactions, or read mapping. The relative scores of different pairs strongly depend on the choice of a substitution matrix and gap penalties, but the existing approaches for the estimation of these parameters do not directly optimize them for the task of classification. In this work, we present DiscrimAlign, a statistical model for discriminative learning of substitution matrices and gap penalties from a dataset of positive and negative pairs of unaligned biological sequences. The model links the alignment score of a sequence pair with the associated binary label through a logistic function and learns the parameters by likelihood maximization. We analyze theoretical properties of the model, derive and implement a learning procedure, study its performance in simulated experiments, and apply it to predict microRNA-target interactions. We show that sequence alignment with discriminative substitution matrices and gap penalties predicts the interactions comparably to state-of-the-art neural network classifiers while being more interpretable. An implementation of the model and reproducibility workflows are available at https://github.com/BioGeMT/DiscrimAlign.

## 1 Introduction

Any kind of alignment of biological sequences requires the user to specify a “substitution matrix” and penalties for gap opening and extension. The quality of the resulting alignment and its usefulness for any given task depends greatly on the choice of these parameters [2, 36]. Alignment of DNA sequences can often be performed with a simple four-parameter scoring scheme, with one score for a match, another for a mismatch, and two penalties for gap openings and extensions. For example, the default option in the BLASTN suite is to assign one point to a match and minus two to a mismatch, although accounting for differences in rates of different mutations has been reported to improve the performance [53]. For protein sequences, more complex scoring schemes are needed to account for similarities of amino acids, such as the PAM and BLOSUM series of substitution matrices. In particular, the BLOSUM62 matrix is the *de facto* standard choice [49, 55, 28, 46], at least in part due to being the default option in BLASTP [6, 52], MUSCLE [14], MAFFT [33], MMseqs2 [54] and others.Despite its popularity, it has been reported that, for some protein families, the pairwise alignment with BLOSUM62 aligns less than 50% of residues correctly [36, 28]. Accordingly, numerous alternative substitution matrices have been developed, both general-purpose and application-specific, both for amino acid and nucleotide sequences [59].

There are several approaches to estimation of substitution matrices, with different underlying “target functions” and intended applications. Most substitution matrices describe patterns of substitutions observed in a collection of ground-truth alignments. Other approaches optimize the similarity between sequence alignment and structural alignment of proteins (see the Related work section). Gap penalties are often hand-selected, but can be also fine-tuned to optimize ortholog prediction or alignment accuracy. Despite the variety of goals for estimation, to our knowledge, there is no approach specifically designed for a binary classification of pairs of sequences. Binary classification covers multiple applications of sequence alignment, including homology search, in which sequences are classified as either related or not; Prediction of protein or RNA interactions; Read mapping, in which a read can be classified as correctly or incorrectly placed on the genome; Detection of duplicated sequences and sequence clustering; Motif or domain searches; and others. These tasks are typically done using empirical substitution matrices and subsequently fine-tuned gap penalties rather than parameters obtained specifically for classification.

### Our contribution

In this work, we explore the task of discriminative learning of substitution matrices and gap penalties for binary classification of pairs of sequences. We consider a data set 𝒟= (*A*_*i*_, *B*_*i*_, *y*_*i*_): *i* = {1, 2, …, |𝒟| *}*, where *A*_*i*_ and *B*_*i*_ are sequences over a common alphabet (typically either DNA, RNA, or amino acid) and *y*_*i*_ ∈ *{*0, 1} is a binary label associated with each pair. Our goal is to design a supervised learning procedure that can distinguish between positive pairs (*y*_*i*_ = 1) and negative pairs (*y*_*i*_ = 0) using alignment score. We analyze a logistic link between the sequences and the label,

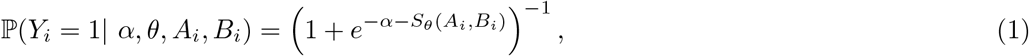

where *θ* denotes a vector of alignment parameters consisting of substitution weights and gap penalties, *S*_*θ*_(*A, B*) denotes the alignment score between sequences *A* and *B* under parameters *θ*, and *α* is an intercept parameter that controls the probability of the pair being positive when the alignment score is zero (corresponding to an empty alignment when the sequences are aligned locally).

The idea to use a logistic function for classification of sequences is well-established and can be derived from the log-odds formalism of substitution matrices [12]. However, the established approach is to use this function to calculate the probability of a relationship between sequences after the alignment score has been calculated under a fixed choice of the substitution matrix and the gap penalties. In this work, we take a reverse approach: we derive an optimization procedure that, given a dataset of positive and negative examples of unaligned sequences, uses the logistic link to infer the parameters. This allows us to learn parameters without example alignments.

Our approach covers the most common types of alignment of sequences over general alphabets, including local, global and semi-global (glocal) alignment schemes, linear and affine gap penalties, symmetric and asymmetric substitution matrices. We present a derivation of the optimization procedure, analyze the properties of the proposed method theoretically and in computational simulation experiments, and present a case study of the prediction of microRNA-target interactions, where we show that the proposed method performs comparably to state-of-the-art neural network classifiers while retaining the interpretability of sequence alignment. A Python implementation of our method is available at https://github.com/BioGeMT/DiscrimAlign, together with Jupyter notebooks with simulation experiments and Python scripts with the case study.

## 2 Related work

The two main approaches to developing amino acid substitution matrices can be described as alignment-based and phylogeny-based. The alignment-based matrices, such as the original BLOSUM [23] and the bug-fixed RBLOSUM and CorBLOSUM series [55, 24, 17], are obtained by counting substitutions in a collection of ground-truth alignments, typically obtained from conserved blocks or by aligning protein structures. This approach has recently been used to obtain the CBM and CCF matrices for *Plasmodium* species [4], the MOLLI series for *Mollicutes* bacteria [39], or the RHOD matrix for microbial rhodopsins [44]. Some alignment-based approaches combine the observed substitutions with other information, for example structural features, which can also account for correlated mutations [58, 28, 35].

While the alignment-based matrices describe patterns observed in a collection of alignments, the phylogeny-based matrices describe patterns of ancestral substitutions inferred from phylogenetic trees. These include the PAM series [11] and the more recent JTT [31], WAG [60], LG [38] and VTML [43, 42] matrices. They are commonly used in the Maximum Likelihood inference of phylogenetic trees, where they are used as matrices of instantaneous rates of substitutions. They can also be used for sequence alignment, possibly after a transformation such as taking a matrix exponent [60, 13]. The phylogenetic approaches seem to focus on general-purpose rather than application-specific matrices, likely due to a considerably more involved estimation procedure [59].

Apart from amino acid matrices, researchers also investigated scoring schemes for nucleotide sequences for homology search [53] and read mapping to reference genomes [16, 21]. Here, multiple sequence or structural alignments are often not available, necessitating a different approach. For example, LAST-TRAIN [21] uses a procedure of iterative realignment of pairs of sequences and recalculation of the substitution matrix based on a Markov Chain formalism of sequence alignment [12, 15]. A similar approach has been used for disordered proteins [5, 50, 41]. Other applications also require substitution matrices obtained from pairs of sequences, such as predicting protein-protein interactions [27], microRNA-target interactions [57], or DNA-to-protein alignment [62].

The selection of appropriate gap penalties for a given substitution matrix is often considered a separate task. The gap penalties for MOLLI matrices were optimized for ortholog prediction [39]. The POP procedure was developed to optimize the accuracy of alignments on a benchmark of structural alignments [13]. A notable exception is the Inverse Parametric Alignment (IPA) approach, which identifies parameters under which a given collection of example alignments has optimal scores, and can thus estimate the substitution matrix jointly with gap penalties [36].

While the approaches described above are often used interchangeably, they have different goals and purposes. Phylogeny-based matrices aim to describe the evolutionary process of substitutions, empirical log-odds matrices summarize patterns observed in a collection of ground-truth alignments, while IPA and POP optimize the similarity of structure-based and sequence-based alignments. While some theoretical guarantees exist for log-odds matrices applied to homology search with an ungapped local alignment [32], overall there is no guarantee that, for example, matrices that describe ancestral substitutions, combined with gap penalties that optimize the structural accuracy of an alignment, will perform optimally when applied to the task of homology search with a local alignment. These considerations highlight the need for developing consistent approaches to parameter estimation, specifically designed for clear end goals.

## 3 Methods

### 3.1 Mathematical properties of pairwise alignment score

In this work, we focus on pairwise sequence alignments that optimize a linear combination of parameters, denoted *θ*, and counts of associated features such as substitutions or gaps, denoted *x*. In the simplest setting, *θ* can be a vector of match, mismatch, and linear gap weights, in which case *θ* = (*w*_match_, *w*_mismatch_, *w*_gap_); the associated configuration vector is then *x* = (*x*_match_, *x*_mismatch_, *x*_gap_), with the numbers of matches, mismatches and gaps in some given alignment. In a setting more commonly used for protein sequences, *θ* consists of either 190 or 400 substitution weights (depending on whether the substitution matrix is assumed to be symmetric) and two gap penalties, and the alignment configuration *x* counts the associated substitutions, gap openings, and gap extensions.

Let *Θ* denote the space of possible parameters and let 𝒳 (*A, B*) denote the set of all alignment configurations that can be obtained from *A* and *B* by inserting any number of gaps, with the exclusion of alignment columns consisting of two gaps only. As the scoring scheme influences the shape of the parameter space *Θ*, the alignment scheme (local, global or glocal) influences the space 𝒳 (*A, B*). For example, the zero vector corresponding to an empty alignment always belongs to the space of configurations of a local alignment, but not of the global one (and, in general, 𝒳_global_(*A, B*) is a proper subset of 𝒳_local_(*A, B*)).

The specification of *Θ* and 𝒳 (*A, B*) is equivalent to the specification of alignment and scoring schemes. In this work, we will assume that *Θ* =ℝ^*n*^ for some *n*; consequently, we will consider an unconstrained optimization problem of parameter learning. Let ⟨*x, y*⟩= _*i*_ *x*_*i*_*y*_*i*_ denote the inner product. Given a fixed configuration space 𝒳 (*A, B*) and some *θ* ∈ *Θ*, the alignment score of sequences *A, B* can be defined as

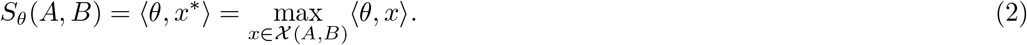

This kind of formulation has previously been used in research on Parametric and Inverse-Parametric Alignment [34, 36, 29]. In general, the space𝒳(*A, B*) has a complex structure and its size grows exponentially with the length of the sequences [19]. The classical alignment algorithm based on dynamic programming serves as a so-called *oracle* that can return an optimal configuration in a polynomial time [36]. We remark that multiple alignments can have the same configuration, and multiple configurations can potentially yield the same optimal score, so *x*^*^ in the definition above denotes one of the optimal configurations that may itself correspond to several alignments.

The formulation of pairwise alignment with Equation (2) allows us to state the following Proposition, which describes the shape of regions of optimality for a given configuration. A set *U* is *convex* if *x, y* ∈ *U* implies *px* + (1 − *p*)*y* ∈ *U* for all *p* ∈ [0, 1]; a set *U* is a *cone* if *x* ∈ *U* implies *sx* ∈ *U* for all *s >* 0.

#### Proposition 1

*Let U*_*A,B*_(*x*) ⊆ *Θ denote the subset of the parameter space on which the alignment configuration x is optimal, and let* 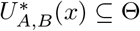*denote the subset in which x is uniquely optimal. Then, U*_*A,B*_(*x*) *is a non-empty closed convex cone, and* 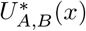*is an open convex cone*.

This result was shown previously in the context of parametric alignment [45] and generalizes previous results about the decomposition of the parameter space, which typically considered some of the parameters to be constant [56, 20]. The proof is based on comparing *x* to every other configuration and expressing *U*_*A,B*_(*x*) as an intersection of the resulting hyperplanes (see the Appendix for details). The cone property is a mathematical expression of the fact that scaling all parameters by a positive constant does not change the relative scores of different alignments, thus preserving the optimal one [2]. The next Proposition describes the dependence of *S*_*θ*_(*A, B*) on *θ*, and will later be necessary to derive generalized gradients for our optimization procedure. A function *f* : ℝ^*n*^ → ℝis said to be Lipschitz continuous with a constant *L* if for any *x, y* ∈ ℝ^*n*^ we have |*f*(*x*) − *f*(*y*)| ≤ *L*||*x* − *y*|| (i.e. the growth of *f* is linearly bounded by *L*). A function *f* is said to be convex if, for any *x, y* ∈ ℝ^*n*^ and *p* ∈ [0, 1], we have *f*(*px* + (1 − *p*)*y*) ≤ *pf*(*x*) + (1 − *p*)*f*(*y*).

#### Proposition 2

*The function* 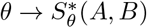 *is convex and Lipschitz continuous with the Lipschitz constant* max_*x*∈*𝒳* (*A,B*)_ ||*x*||.

Both properties stem from the fact that for any vector *x*, the scalar product ⟨*θ, x*⟩ is a convex and Lipschitz continuous function of *θ*, and the maximum operator over a finite number of functions preserves these properties (see the Appendix for a detailed proof). The Rademacher theorem states that Lipschitz-continous functions are differentiable almost everywhere (i.e. the set of points of non-differentiability has a Lebesgue measure zero). In the case of *S*_*θ*_(*A, B*), if *θ*∈*U*^*^(*x*) for some *x*, then ∇_*θ*_*S*_*θ*_(*A, B*) exists at *θ* and is equal to *x*. On the other hand, if two or more different configurations are optimal at *θ*, then *S*_*θ*_(*A, B*) is non-differentiable at *θ*. It follows that the optimal configurations are unique for almost all values of *θ*.

The non-existence of gradients of *S*_*A,B*_(*θ*) at some points makes standard optimization procedures like gradient descent not applicable for parameter learning. This problem has recently been overcome by designing differentiable alignment algorithms [48]. However, since *S*_*A,B*_(*θ*) is convex, it has a *subdifferential*, which is a generalization of a gradient for convex functions and will allow us to develop learning procedures for the standard, non-differentiable alignment algorithms.

#### Definition 3

*(Adapted from [3]) A subdifferential of a convex function f* : ℝ^*n*^ →ℝ *at a point x, denoted* ∂*f*(*x*), *is the set of coefficient vectors of linear functions which bound f from below (also referred to as supporting hyperplanes of f at x):*

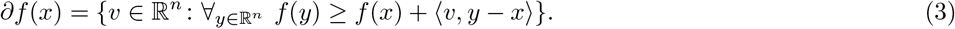

*A subgradient of f at x is any element of the subdifferential* ∂*f*(*x*).

For example, if *f*(*x*) =|*x*|, then ∂*f*(*x*) = {−1} for all *x <* 0, ∂*f*(*x*) = {1} for all *x >* 0, and ∂*f*(0) = [ −1, 1]. Similarly to standard gradients, subgradients indicate the direction of growth of the function: for any *v* ∈∂*f*(*x*), setting *y* = *x* + *λv* in Equation (3) results in *f*(*x* + *λv*) −*f*(*x*) ≥⟨*v, λv*⟩ = *λ*|| *v*|| ^2^, indicating that *f*(*x* + *λv*) ≥*f*(*x*) whenever *λ*≥ 0. If follows that *x* is the minimum of *f*(*x*) if and only if 0∈∂*f*(*x*) [3]. However, unlike standard gradients of differentiable functions, the definition does not guarantee that −*v* indicates the direction in which the function decreases.

Let Conv(*X*) denote the convex hull of a set *X*, i.e. the minimal convex set containing all points from *X* (equivalently, the set of all convex combinations of points from *X*). For a function *f*(*x*) = max_*i*∈*I*_ *f*_*i*_(*x*), if *I* is a finite set and *f*_*i*_ are convex, then ∂*f*(*x*) = Conv U_*i*_*{*∂*f*_*i*_(*x*): *f*_*i*_(*x*) = *f*(*x*)*}* (see e.g. Theorem 3.50 in ref. [3]).

#### Proposition 4

*Let S*_*θ*_(*A, B*) *be the optimal score of alignment of sequences A and B with parameter vector θ. Then, the subdifferential of the function θ* →*S*_*A,B*_(*θ*) *is the convex hull of all configurations optimal at θ:*

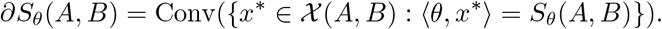

From the Proposition it follows that any optimal configuration is a subgradient of *S*_*θ*_(*A, B*), and can be obtained by calculating the score with the dynamic programming algorithm, back-tracking the optimal alignments, and calculating the configurations. If only one subgradient is necessary for any given *θ*, it can be calculated without backtracking by simply keeping track of the choices during the calculation of the score with the dynamic programming algorithm.

### 3.2 The likelihood function of the proposed model

Let *p*(*α, θ*; *A*_*i*_, *B*_*i*_) = ℙ(*Y*_*i*_ = 1 |*θ, α, A*_*i*_, *B*_*i*_). In the same way as in logistic regression models, we assume that the likelihood of parameters *α* and *θ* given a dataset 𝒟is equal to the product of probabilities defined by Equation (1):

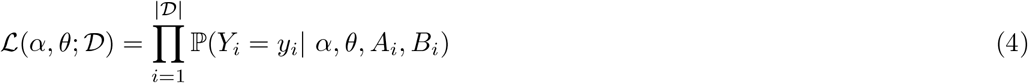

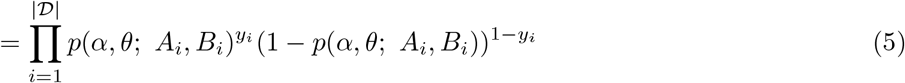

Given a dataset 𝒟, the parameters *α* and *θ* can be estimated by maximizing the above likelihood function. This is equivalent to maximizing the log-likelihood function *l*(*α, θ*;𝒟 ) = ln ℒ (*α, θ*; 𝒟 ). Let 𝒳_*i*_ = 𝒳 (*A*_*i*_, *B*_*i*_). After taking the logarithm of Equation (5) and some algebraic manipulations, the log-likelihood of the proposed model can be expressed as

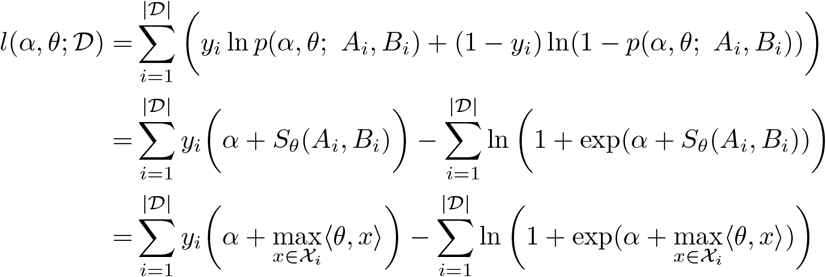

The main difference between the proposed model and the standard logistic regression is the presence of the maximum operator within the log-likelihood. The model reduces to the standard logistic regression when conditioned on alignments. Namely, consider a list of fixed (possibly sub-optimal) alignments 𝒜_*i*_ = 𝒜 (*A*_*i*_, *B*_*i*_). Given this list, the probability of a pair being positive can then be expressed as

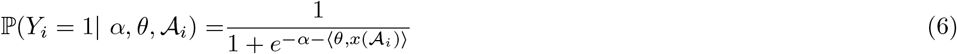

This equation corresponds to a standard logistic regression model with intercept *α*, vector of parameters *θ*, and independent variables *x*(𝒜 _*i*_). If a list of ground-truth alignments is available, this model can be used to estimate the parameters, but does not guarantee that the alignments will be optimal for parameters estimated this way (especially if the alignments are inaccurate). The model proposed in Equation (5) uses unaligned pairs of sequences and estimates the alignments jointly with the parameters, allowing both to change during optimization.

Note that functions *y*_*i*_(*α* + *S*_*θ*_(*A*_*i*_, *B*_*i*_)) are convex as affine transformations of a convex function *S*_*θ*_(*A*_*i*_, *B*_*i*_)), functions ln (1 + exp(*α* + *S*_*θ*_(*A*_*i*_, *B*_*i*_))) are convex as compositions of a strictly increasing convex function ln(1 + exp(*α* + *x*)) and convex functions *S*_*θ*_(*A*_*i*_, *B*_*i*_), and convexity is preserved under summation. The log-likelihood of the proposed model is therefore a difference of two convex functions, and such a difference may be neither convex nor concave (as opposed to the log-likelihood of the standard logistic regression, which is concave). However, it is concave locally, meaning that for any *x*∈ ℝ^*n*^ there exists an open neighborhood *U*∋*x* such that the function restricted to *U* is concave (i.e. −*l*(*α, θ*;𝒟 ) _|*θ*∈*U*_ is convex).

#### Proposition 5

*The function l*(*α, θ*;𝒟 ) *is smooth and locally concave with respect to θ almost everywhere*.

The Proposition stems from the fact that the log-likelihood is differentiable almost everywhere, and in the points of differentiability it locally reduces to the log-likelihood of a standard logistic regression. A detailed proof is presented in the Appendix. Since *l*(*α, θ*; 𝒟) is not globally concave, the model may not be uniquely identifiable (i.e. there may be more than one maximizer of the log-likelihood). An intuitive explanation of this phenomenon is that the added flexibility of alignment means that different substitution matrices may perform equally well in classifying pairs of sequences, as the differences in substitution matrices can potentially be offset by adjusting the alignments. As a consequence, we will look for example maximizers *α*^*^, *θ*^*^ ∈ arg max_*α,θ*_ℒ (*α, θ*; 𝒟).

#### Proposition 6

*The set of maximizers α*^*^, *θ*^*^∈arg max_*α,θ*_ℒ (*α, θ*; 𝒟) *has Lebesgue measure zero*.

The Proposition follows from the fact that *Θ* is composed of a finite number of open subsets on which↕(*α, θ*; 𝒟) is concave (and thus has a unique maximizer), and the boundaries of these subsets, which have measure zero. An exact characterization of the set of maximizers of Equation (5) remains an open problem, in particular its convexity and conditions for its boundedness. A necessary condition for the latter is the lack of a complete separation of the labels (i.e. the possibility to perfectly predict them in the training set), similarly as in the standard logistic regression [1]:

#### Proposition 7

*Assume there exists a* threshold *c* ∈ℝ *and a parameter vector θ such that S*_*θ*_(*A*_*i*_, *B*_*i*_) *> c for any i such that y*_*i*_ = 1 *and S*_*θ*_(*A*_*i*_, *B*_*i*_) *< c for any i such that y*_*i*_ = 0. *Then*, sup_*α,θ*_ ℒ (*α, θ*; 𝒟) = 1. *Furthermore, for any sequence α*_*k*_, *θ*_*k*_ *such that* lim_*k*→∞_ ℒ (*α*_*k*_, *θ*_*k*_; 𝒟) = 1, *the alignment parameters diverge to infinity:* ||*θ*_*k*_|| → ∞.

**Proof**. Let *α*_*k*_ = −*kc, θ*_*k*_ = *kθ* and observe that *S*_*θk*_ (*A*_*i*_, *B*_*i*_) = *kS*_*θ*_(*A*_*i*_, *B*_*i*_). We have

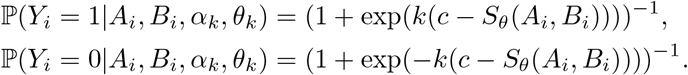

Since *c*− *S*_*θ*_(*A*_*i*_, *B*_*i*_) *<* 0 for *y*_*i*_ = 1 and *c* −*S*_*θ*_(*A*_*i*_, *B*_*i*_) *>* 0 for *y*_*i*_ = 0, for every *i* we have ℙ(*Y*_*i*_ = *y*_*i*_ | *A*_*i*_, *B*_*i*_, *α*_*k*_, *θ*_*k*_) →1 as *k*→ ∞, which proves the first part of the Proposition. The second part follows from the fact that the likelihood function attains 1 only if all the logistic probabilities attain 1, which can only happen in the limit of infinite alignment scores. ◀

We also conjecture, motivated by similar results in standard logistic regression [1] and the results of our numerical experiments, that a so-called overlap of points (i.e. a dataset in which any set of parameters results in at least one misclassified label) is sufficient for there to exist finite maximizers *α*^*^, *θ*^*^ (and, possibly, for the set of maximizers to be bounded). In what follows, we will assume that the set of maximizers of *l*(*α, θ*; 𝒟 ) for a given dataset 𝒟is non-empty.

### 3.3 The Clarke subdifferential of the log-likelihood

As opposed to the function *θ*→*S*_*θ*_(*A, B*), the functions ln *p*(*θ*) and ln(1−*p*(*θ*)) may not be convex nor concave. This is because the logarithm of the logistic function is concave, while the alignment score is convex, and a composition of a concave and a convex function may be neither. Thus, these functions may not have standard subdifferentials, and in order to optimize the log-likelihood we need to use a further generalization called the Clarke subdifferentials [7, 8]. A function is said to be locally Lipschitz continuous if, for every *x* ∈ ℝ^*n*^, there exists an open neighbourhood *U* ∋ *x* such that *f* is Lipschitz continuous on *U*.

#### Definition 8

*(Adapted from [7]) For a locally Lipschitz function f* : ℝ^*n*^→ ℝ, *the Clarke subdifferential at a point x, denoted* ∂^*C*^*f*(*x*), *is defined as:*

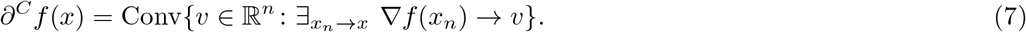

Intuitively, one may think about the Clarke subdifferential as the set of all gradients of *f*, and their convex combinations, around *x*. This intuition is substantiated by the fact that locally Lipschitz functions are differentiable almost everywhere, so their gradients exist almost everywhere. Points *x* such that 0∈∂^*C*^*f*(*x*) are called *Clarke-stationary* or *Clarke-critical*. Similarly as in standard calculus, if *f* has a local minimum or maximum at *x*, then *x* is a Clarke-stationary point, but the converse may not hold (Proposition 2.3.2 in [8]). As opposed to standard calculus, the rules of Clarke calculus often provide only inclusions rather than equalities.

#### Lemma 9

*(Consequence of Theorem 2*.*3*.*9 in [8]). Let f, g* : ℝ^*n*^ → ℝ *be locally Lipschitz functions, and let h*: ℝ → ℝ *be locally Lipschitz and continuously differentiable (C*^1^*). Let* ⊕*denote the Minkovsky sum of sets defined as A* ⊕ *B* = *{a* + *b* : *a* ∈ *A, b* ∈ *B}. Then*,

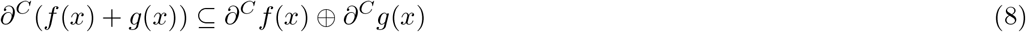

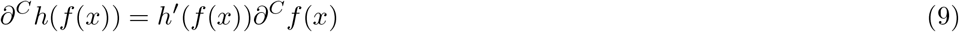

We remark that the chain rule above works only in the special case that *h* is a continuously differentiable function of a single variable; in general, only an inclusive chain rule is satisfied with ∂^*C*^*h*(*f*(*x*))⊆ *h*^*′*^(*f*(*x*))∂^*C*^*f*(*x*). In what follows, we will denote the Clarke subdifferential with respect to *θ* as 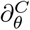, treating the parameter *α* as a constant.

#### Proposition 10

*The Clarke subdifferentials of* ln *p*(*α, θ*; *A, B*) *and* ln(1 − *p*(*α, θ*; *A, B*)) *with respect to θ are equal to*

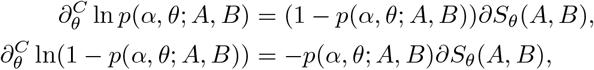

The proof is based on showing that the functions are locally Lipschitz continuous and applying Lemma 9. Details are presented in the Appendix. Using a variable *y*∈{ 0, 1}, we can combine both equations from the Proposition into

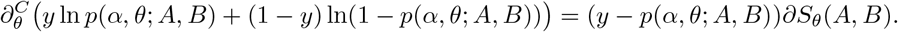

This equality follows from the fact that any choice of *y*∈ {0, 1} reduces this equation to either of the two from the Proposition. From Lemma 9 we now have

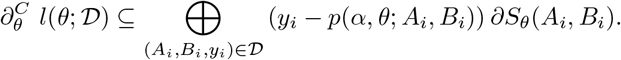

In general, 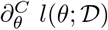 is a proper subset of the Minkovsky sum on the right-hand side. This stems from the fact that the Minkovsky sum allows us to select any optimal configuration for each sequence pair, while the Clarke subdifferential requires a single convergent sequence of parameters *θ*_*n*_ → *θ* for which the configurations are jointly optimal:

#### Lemma 11

*Let* ℬ (*θ*; 𝒟) = *{*(*x*_1_, *x*_2_, …, *x*_|𝒟|_): *x*_*i*_ ∈ ∂_*θ*_*S*(*θ*; *A*_*i*_, *B*_*i*_)*}. Then*,

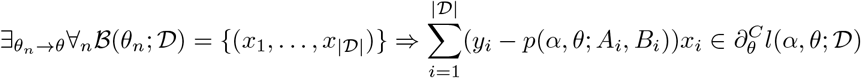

A proof of the Lemma is presented in the Appendix. The Lemma can be used to show the following Proposition. We remark that both the Lemma and the Proposition state an inclusion rather than equality due to the technical fact that the sum of non-differentiable functions may be differentiable. This issue is discussed further in the proof of the Lemma.

#### Proposition 12

*Let* (*x*_1_, *x*_2_, …, *x* _*|𝒟|*_) ∈ ℬ (*θ*; 𝒟). *For a configuration x*_*i*_ ∈𝒳_*i*_, *let* 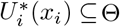*be the set of parameters θ for which x*_*i*_ *is a uniquely optimal alignment configuration of sequence pair A*_*i*_, *B*_*i*_. *Then, if the intersection of the sets* 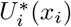*is non-empty, then*

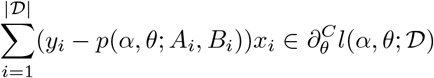

The subgradients of *l*(*α, θ*; 𝒟) above have an intuitive interpretation. The weights *y*_*i*_ − *p*(*α, θ*; *A*_*i*_, *B*_*i*_) are the errors of the model, which are positive for the positive pairs and negative for the negative ones. Thus, they prioritize increasing the parameters corresponding to common substitutions in low-scoring positive pairs and decreasing the ones corresponding to common substitutions in high-scoring negative pairs. We also note a connection to the problem of separability: if there exists a sequence *θ*_*k*_ such that lim_*k*→∞_ *p*(*α, θ*_*k*_; *A*_*i*_, *B*_*i*_) = *y*_*i*_ for all *i*, then 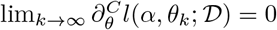, resulting in “Clarke-stationarity in infinity”.

We remark that functions whose gradients point towards the direction of increase are termed *pseudoconvex*. The following Proposition establishes that ln *p*(*α, θ*; *A, B*) is a Clarke-pseudoconvex function and ln(1 −*p*(*α, θ*; *A, B*)) is a Clarke-pseudoconcave function. Whether 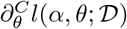admits a similar property remains an open question.

#### Proposition 13

*Let* 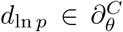ln *p*(*α, θ*; *A, B*) *and* 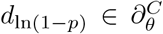ln(1 − *p*(*α, θ*; *A, B*)).

*Then, for any θ*^*′*^ ∈ *Θ:*

▄ *If* ⟨*d*_ln *p*_, *θ*^*′*^− *θ*⟩≥ 0, *then* ln *p*(*α, θ*^*′*^; *A, B*) ≥ln *p*(*α, θ*; *A, B*),
▄ *If* ⟨*d*_ln(1 − *p*)_, *θ*^*′*^− *θ*⟩≤ 0, *then* ln(1− *p*(*α, θ*^*′*^; *A, B*)) ≤ln(1− *p*(*α, θ*; *A, B*)).

The proof is based on an observation that *d*_ln *p*_ is proportional to a subgradient of the

alignment score, and thus points in the direction of increasing alignment score, which in turn is the direction of increase of ln *p*(*α, θ*; *A, B*) for a fixed *α*.

### 3.4 Fitting the model with the subgradient method

While there are specialized methods of optimizing differences of convex functions such as the log-likelihood of the proposed model [25, 30], in this work we present a general subgradient method because it results in a particularly simple algorithm that can be implemented in a few lines of computer code. The subgradient method is a generalization of gradient descent that essentially replaces the gradient with any subgradient. While the definition of subgradients does not guarantee that the function value increases in each step (hence the name “subgradient method” rather than “subgradient ascent”), the method converges to a minimizer under some general assumptions. We will first present the conditions for finding optimal *θ* assuming that the *α* parameter is known. Below, “almost surely” means “with probability 1” and refers to the randomness of the trajectory.

#### Theorem 14

*For k* ≥1, *let ξ*_*k*_ *be an n-element vector of independent random variables with zero expected value and standard deviation σ*_*k*_, *where n is the dimension of Θ. Let θ*_0_ ∈ *Θ and let the sequence θ*_*k*_ *be defined as*

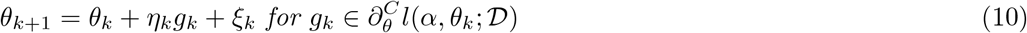

*Assume that* 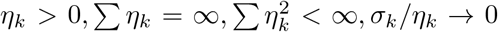*and assume that the sequence θ*_*k*_ *is bounded. Then, almost surely, every limit point of θ*_*k*_ *is a Clarke-stationary point of l*(*α, θ*; 𝒟).

The proof is based on showing that the sequence of parameters satisfies Assumptions C and D from ref. [10], and presented in the Appendix. The lack of complete separability is a necessary condition for the existence of finite global maximizers, but does not preclude the existence of local maxima or other Clarke-stationary points. Sufficient conditions for the existence of stationary points, maximizers, and the boundedness of ||*θ*_*k*_|| remain an open question. Convergence issues can be detected in practice by monitoring the trajectory of||*θ*_*k*_|| to detect unboundedness.

An example sequence of step-lengths that satisfies the assumptions of the Theorem is *η*_*k*_ = *η/k* for some *η >* 0 and *σ*_*k*_ = *σ/k*^2^ for some *σ* ≥0. Note that while *σ*_*k*_≡ 0 also satisfies the assumptions of the Theorem, some amount of random noise may be beneficial in practice e.g. to avoid local stationary points. We also remark that the assumption *σ*_*k*_*/η*_*k*_→0 can be considerably weakened to still preserve convergence, but having the random noise vanish faster than the step size allows the method to refine the estimates without random perturbations in final iterations. Finally, while subgradients of the log-likelihood can be theoretically constructed using Proposition 12, it might be computationally expensive in practice. However, if *ξ*_*k*_ are continuous random variables, then for *k* ≥ 1 the log-likelihood is differentiable at *θ*_*k*_ almost surely. This reduces the condition 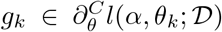to 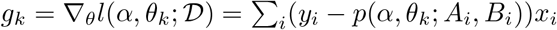 almost surely.

The downside of Theorem 14 is that it requires that all alignments be recalculated at each step, making it computationally intensive for large datasets. This can be mitigated by taking a random sample of sequence pairs at each iteration (similar to a batch learning strategy in neural networks). In the Theorem below, subgradients are calculated using random samples 𝒟_*k*_, but the parameters converge to a stationary point corresponding to the full dataset𝒟 .

The proof is presented in the Appendix.

#### Theorem 15

*For k* ≥1 *and N* ∈ ℕ_+_, *let 𝒟_k_ be a sequence of independent N -element random subsets of 𝒟 such that the elements of each subset are selected uniformly with replacement. Assume the same conditions on η_k_ and ξ_k_ as in Theorem 14, but let the sequence θ_k_ now be defined as*

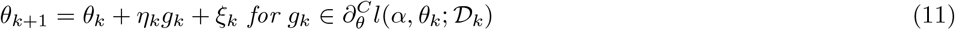

*Assume that for each k, subgradient g*_*k*_ *is obtained using Proposition* (12) *applied to l*(*α, θ*_*k*_; **𝒟**_*k*_). *Then, with probability 1, every limit point of θ*_*k*_ *is a Clarke-stationary point of l*(*α, θ*; **𝒟**).

Theorems 14 and 15 provide ways of estimating *θ* under a known *α*. In order to optimize the latter, we note that the log-likelihood is differentiable and concave (due to a negative second derivative) with respect to *α* for any fixed *θ*. It follows that for any fixed *θ* the optimal value of *α* is unique and can be efficiently found with standard optimization methods such as the Newton-Rhapson procedure. We use this observation to replace the free parameter *α* with *α*^*^(*θ*) = arg max_*α*_ *l*(*α, θ*; **𝒟**) and optimize *l*(*α*^*^(*θ*), *θ*; **𝒟**) using the subgradient method for *θ*. This procedure is summarized in Algorithm 1.

The downside of subgradient methods is their slow rate of convergence and the lack of simple stopping criteria. A common practice is to run the method for a given number of iterations and select the best value from the whole trajectory (not necessary at the final iteration). However, in our experiments, we see that the trajectory of the log-likelihood generally stabilizes after some number of iterations, and we simply select the final value. While theoretical guarantees of convergence require a decreasing sequence of steps, in practice we generally see a better performance with a constant step size *η*_*k*_≡*η* due to a faster convergence, and we use this sequence in the Results section with *σ*_*k*_ = *σ/k*. In order to monitor the convergence, we record the values of *l*(*α*_*k*_, *θ*_*k*_; **𝒟**) and ||*g*_*k*_||.

#### Algorithm 1

Discriminative learning of alignment parameters from pairs of labeled sequences

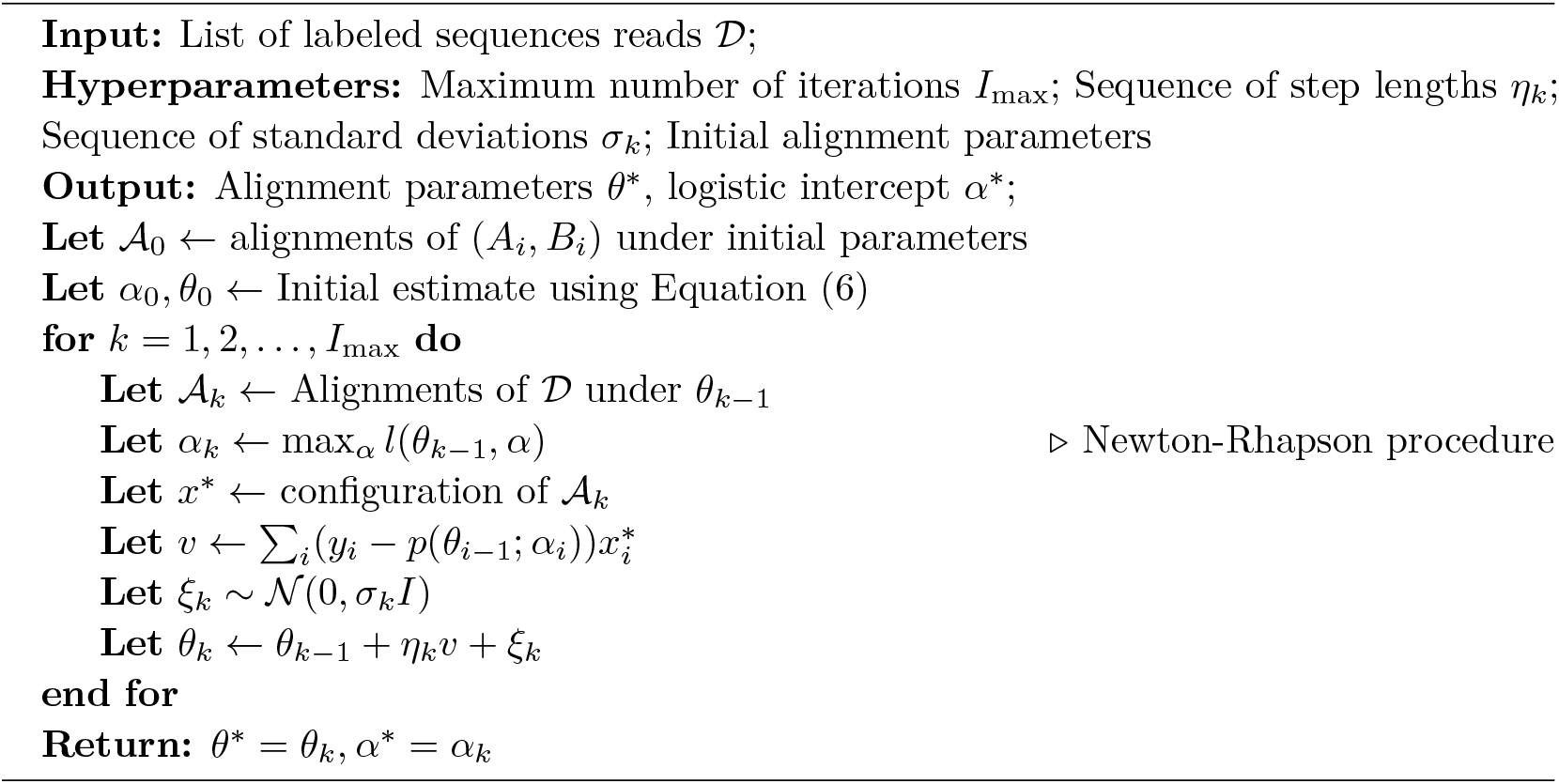

## 4 Results

### 4.1 Convergence and accuracy of estimation

To experimentally validate the proposed method, we performed simulation experiments with fixed substitution matrices and gap penalties using a data set of microRNA (miRNA) sequences. MiRNAs are short RNA molecules (typically 18-25 nt) that regulate gene expression by recognizing mRNA transcripts, or other RNA molecules, through partial complementary pairing and inhibiting translation and/or promoting degradation [40, 37, 26]. Recently, a set of benchmark datasets called miRBench was released, which contains several datasets of pairs of miRNA sequences and 50nt-long RNA sequences labeled based on whether the RNA sequence contains a target site for the miRNA [51, 22]. We used the train split of the Hejret dataset from miRBench, which contains 4084 positive and 4109 negative instances. We ignored the original labels and generated simulated ones in order to obtain a dataset with realistic pairs of sequences and with a known ground truth.

We investigated two local alignment scoring schemes, referred to as “simple model” and “full model”. The “simple” model had match weight 1, mismatch weight -1, gap open -1.2 and gap extend -0.1 The more challenging “full” model had an asymmetric DNA substitution matrix *M* with manually selected entries to obtain a diversity of scores:

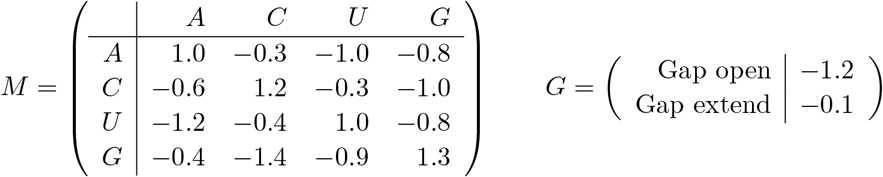

To generate labels, we calculated the scores of pairwise local alignments of miRNA sequences and RNA fragments using Biopython [9]. We used the scores to calculate the probabilities of interactions using Equation (1), with *α* equal to the negative of the median score to obtain a roughly balanced dataset. Finally, for each pair, we used the corresponding probability to simulate a binary label. We run DiscrimAlign on the simulated dataset for 200 iterations with a stochastic factor *σ* = 0.001 and different learning rates *η* ranging from 5e-06 to 5e-05 (using a constant step size in order to obtain a faster convergence, as discussed in Methods).

The results are shown in Figure 1. Step sizes which resulted in an initial drop in the log-likelihood often corresponded to a faster convergence. Excessive step sizes resulted in an unstable behavior, while insufficient step sizes result in a very slow convergence. The step sizes *η* = 2 ·10^−5^ for the simple model and *η* = 4 ·10^−5^ for the full model were among the smallest step sizes that resulted in a sufficiently fast convergence, thus offering a balance between speed and numerical stability. Fig. 2 shows the estimated weights of the asymmetric matrix for *η* = 4 *·* 10^−5^. The correlation between the entries of the true substitution matrix *M* and the estimated one 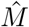 was equal to over 0.99, and the mean absolute error of estimation was 0.06. To analyze the robustness of the method, we run DiscrimAlign with *η* = 4· 10^−5^ for 20 replicated datasets with the asymmetric matrix. The results show that the speed of convergence varies, but the true value of the log-likelihood was reached (or nearly reached) after 200 iterations in all replicates (Fig. 1). The mean absolute error varied between 0.06 and 0.12.

**Figure 1.**
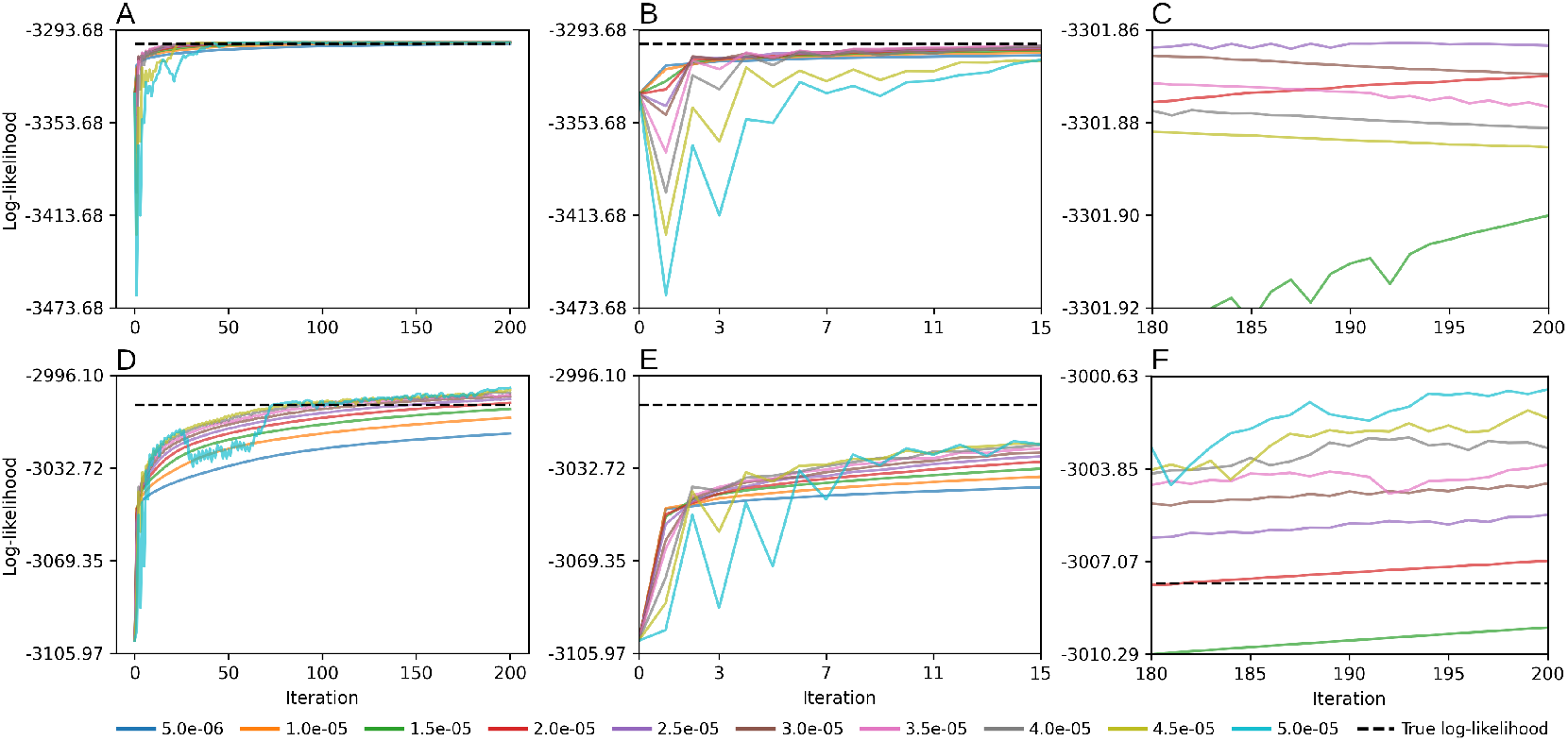
Estimation of discriminative substitution matrices with DiscrimAlign for pairs of RNA sequences with simulated labels. Plots show trajectories of the log-likelihood for different step sizes *η* for a constant sequence of step sizes. A,B,C: Simple substitution model with match/mismatch weights and an affine gap penalty (4 parameters plus intercept). D,E,F: Asymmetric substitution matrix and an affine gap penalty (18 parameters plus intercept). A,D: Full trajectories; B,E: Initial iterations; C,F: Final iterations. All trajectories shown in panel C are above the true log-likelihood; the remaining trajectories did not reach it or were unstable.

**Figure 2.**
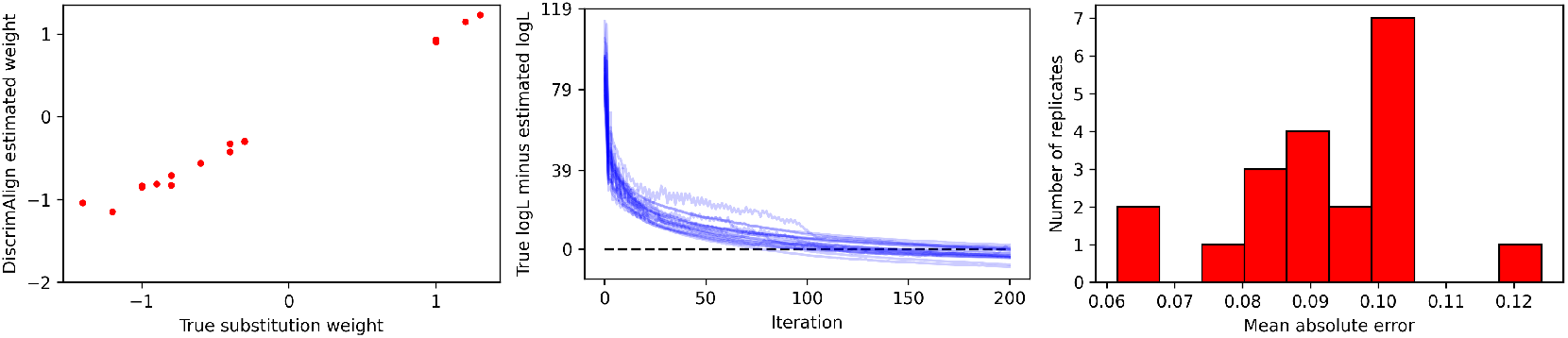
Accuracy of estimation in replicated experiments on RNA sequences with simulated labels. Left: Estimated vs ground-truth substitution weights in an example run. Right: Trajectories of the difference between current log-likelihood and the true log-likelihood for 20 replicated experiments. Right: A histogram of mean absolute errors of estimation for 20 replicated experiments. All plots correspond to a model with an asymmetric substitution matrix and an affine gap penalty.

### 4.2 Dataset requirements for amino acid matrices

A natural question is the size of dataset necessary for an accurate estimation depending on the size of the model and the alphabets. For example, a full model for amino acid sequences would require 403 parameters (400 substitution weights, 2 gap penalties, and the intercept *α*). The error of estimation can be decomposed into error resulting from the training procedure (which includes learning the alignments) and the inherent statistical error resulting from the finite size of the dataset and randomness of labels. In this section, we show how to estimate the latter error in simulated experiments using a standard logistic regression model applied to ground-truth alignments (Equation (6)). This provides a lower bound on the error of DiscrimAlign without the need for a full training procedure, while also allowing one to use an extensive body of statistical results including confidence intervals. In turn, a lower bound on the error provides a lower bound on dataset requirements.

As the ground truth substitution matrix, we used a centered and scaled BLOSUM62 matrix with gap penalties *G* = (−1, −0.1). Scaling the matrix was done so that individual substitutions do not have an overwhelming impact on the predicted probability, as large values of *θ* mean that any differences in alignment configurations result in large changes of the predicted probability. To generate sequence pairs, we downloaded proteomes of *Homo sapiens* and *Gallus gallus* from the NCBI Genome database (proteomes `GCF_000001405.40`and `GCF_016699485.2` respectively). We selected proteins up to 1 000 amino acids in length to avoid excessive computational times, used a local installation of BLASTP to identify homologs with an E-value threshold of 1e-6 (resulting in 2 238 760 pairs), and created three datasets by randomly selecting pairs with |*𝒟*| = 10^4^, 10^5^ or 10^6^. We calculated global alignments and discarded those which consisted of more than 20% of gaps. We used the scores of remaining alignments to generate labels as in the previous section. Finally, we have arranged the configurations of the alignments and the labels into arrays and estimated the parameters (intercept, asymmetric subsitution matrix, and gap penalties) using the LogisticRegression function implemented in the scikit-learn library ( `penalty=‘None’, solver=‘newton-cg’`) [47].

The large dataset |*𝒟*| = 10^6^ resulted in a clear correlation between the estimated and ground-truth substitution weights (Fig 3), providing a likely indication of the minimal requirements for an accurate estimation of discriminative substitution matrices for detecting homology between *Homo sapiens* and *Gallus gallus* proteins. For =|*𝒟*| 10^4^, the estimated parameters were only loosely correlated with the ground-truth, which is to be expected considering the large number of parameters. Our experiments indicated that this correlation depends not just on *N*, but also on the scaling of the substitution matrix; small parameter values were more difficult to identify by logistic regression, because they had little impact on probability. A realistic range of substitution weights for amino acid discriminative substitution matrices requires further studies, and most likely depends on the similarity of compared proteins. The construction of proper datasets for estimation of discriminative matrices for homology search also requires further studies, as selecting positive and negative instances simply as related and unrelated proteins results in a complete separation. This can be overcome either by a specific selection of the dataset or developing regularized models.

**Figure 3.**
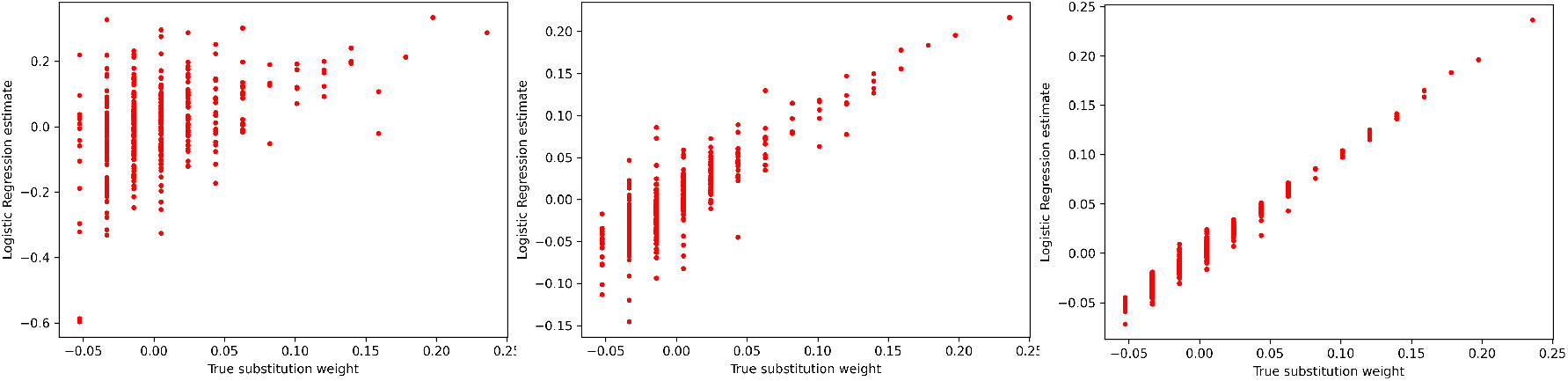
A lower bound on the estimation error for asymmetric amino acid substitution matrices for different dataset sizes. Logistic regression estimation of amino acid substitution matrices directly from optimal alignments obtained with the true parameters (without DiscrimAlign optimization). From left to right, |*𝒟*| = 10^4^, 10^5^, 10^6^.

### 4.3 Case study: prediction of miRNA-target interactions

The task of predicting whether a given pair of miRNA and RNA sequences interact is challenging and typically addressed with black-box methods such as convolutional neural networks [51], although approaches based on alignments with position-specific substitution matrices have been explored with promising results [57]. In this case study, we investigated whether a pairwise alignment score with a discriminative substitution matrix can be used to identify interacting pairs. We used the *D*_Hejret_ dataset and another, larger dataset from the miRBench package [51], referred to as Manakov, with 1 253 320 positive instances in the training split. We run DiscrimAlign on pairs of miRNA sequences and reverse-complemented target sequences with labels indicating experimentally confirmed interactions. We investigated all combinations of linear and affine gap penalties and simple, symmetric and asymmetric substitution matrices. We evaluated the performance using the area under the precision-recall curve (AUPRC) on the test split of each dataset in the same way as reported in the miRBench article [51].

Each training dataset was divided into a fitting subset and a validation subset. The fitting subset was used to estimate the alignment parameters for each candidate model, while the validation subset was used to select the best configuration. The selected configuration was then refitted on the full training split and evaluated on the held-out test datasets. On the Hejret dataset, this resulted in the following parameters for the full model:

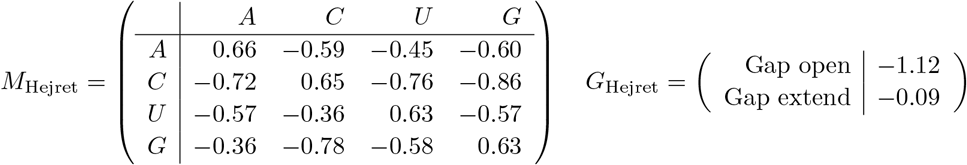

The full models provided the best AUPRC when evaluated on the held-out test splits of their respective datasets, but not in cross-dataset evaluation, indicating some degree of overfitting to the experimental technique used in each dataset (Table 1). Models with a symmetric substitution matrix seemed to generalize better between datasets while having only slightly lower performance within datasets. Overall, models trained on the larger 𝒟_Manakov_ generalized well to the Hejret test set, achieving AUPRC values close to those of the Hejret-trained models, which was not the case for models trained on 𝒟_Hejret_ when evaluated on 𝒟_Manakov_. This suggests that the larger Manakov training set may capture broader miRNA-target pairing patterns. The results of DiscrimAlign were comparable to several state-of-the-art classifiers based on neural networks, including the ones that had the

**Table 1.** The area under precision-recall curve (AUPRC) of miRNA interaction prediction for different alignment models trained with DiscrimAlign.

| Training dataset | Hejret Train |  |  | Manakov Train |  |  |
| --- | --- | --- | --- | --- | --- | --- |
| Test dataset | Hejret Test | Manakov Test | Manakov Leave-Out | Hejret Test | Manakov Test | Manakov Leave-Out |
| Simple Linear | 0.851 | 0.750 | 0.745 | 0.851 | 0.752 | 0.747 |
| Simple Affine | 0.851 | 0.762 | 0.758 | 0.846 | 0.764 | 0.761 |
| Symmetric Linear | 0.859 | 0.757 | 0.752 | **0.853** | 0.760 | 0.759 |
| Symmetric Affine | 0.862 | **0.765** | **0.759** | 0.849 | **0.766** | 0.763 |
| Asymmetric Linear | **0.866** | 0.750 | 0.746 | 0.852 | 0.761 | 0.759 |
| Asymmetric Affine | **0.866** | 0.757 | 0.753 | 0.848 | **0.766** | **0.764** |

best performance reported in the miRBench article (Table 2).

**Table 2.**
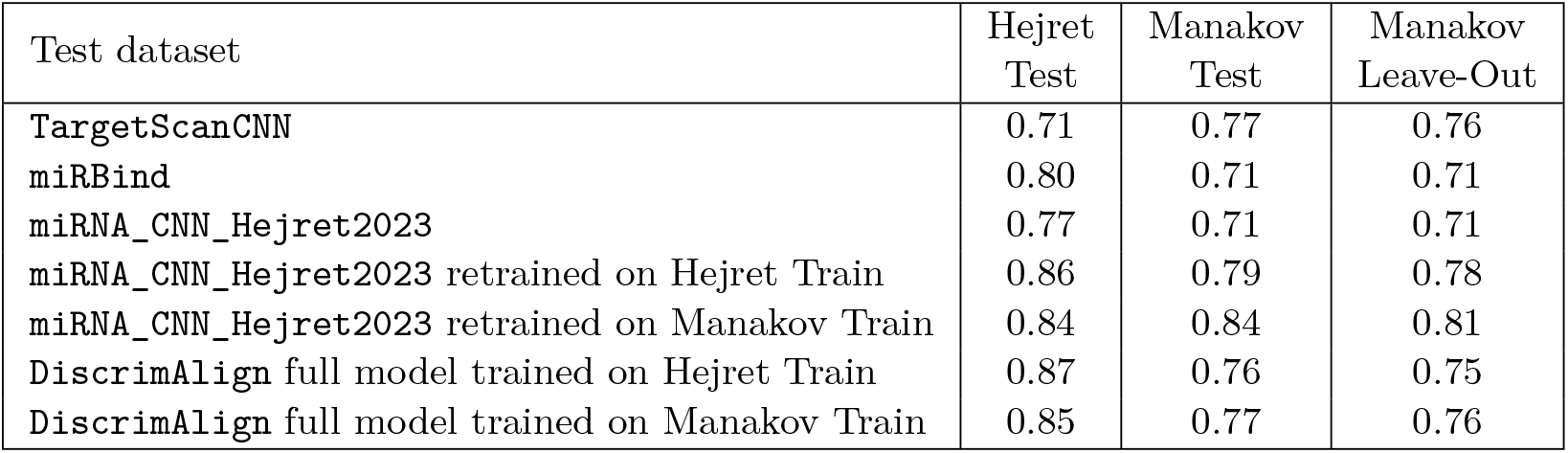
The Area under Precision-Recall curves for different miRNA-target prediction algorithms. Data for `TargetScanCNN, miRBind` and `miRNA_CNN_Hejret2023` comes from ref. [51], in which `miRNA_CNN_Hejret2023` retrained on each dataset was reported to have the highest AUPRC out of the models compared therein.

## 5 Discussion

Some of the first substitution matrices, both for amino acids and nucleotides, were derived from first principles, such as the physical properties of amino acids or the properties of the genetic code [46, 18, 32]. When databases of example alignments became available, researchers started to derive empirical substitution matrices, either from a collection of alignments or from ancestral substitutions inferred from phylogenetic trees. In this work, we propose a new type of substitution matrices that we call *discriminative substitution matrices*. As opposed to empirical matrices which describe patterns observed in related sequences, discriminative matrices describe the differences observed between two sets of pairs of sequences.

We have described a general theoretical framework of learning discriminative matrices through a logistic link between the alignment score and the label associated with a given pair. The framework is based on an analysis of the mathematical properties of alignment score and the logistic link as functions of substitution and gap weights. We have derived a learning procedure based on the subgradient optimization method, which we validated on simulation experiments to show convergence and accuracy of estimation on RNA sequences, and analyzed dataset requirements for aminoacid sequences. Finally, we applied the method to learn a discriminative matrix for the task of predicting microRNA-target interactions. The discriminative alignment performed similarly to state-of-the-art classifiers based on neural networks while offering more biological insight into the nature of the interactions thanks to the interpretable nature of the model, which can be used in a downstream analysis to identify binding patterns by investigating the optimal alignments.

The proposed model offers many topics for future investigations, some of which were mentioned throughout the paper. Other topics include improvements in computational efficiency; Estimating discriminative amino acid substitution matrices on larger datasets; or deriving statistical tests for parameters, such as a likelihood ratio test to determine whether an affine gap penalty gives a statistically significant improvement in classification compared to a linear one in a particular application. Finally, while the proposed method is not explicitly designed to optimize the accuracy of alignments, it would be interesting to see how it performs compared to methods such as IPA or POP.

## Acknowledgements

We thank Mr Dave Galea for the help in running simulations and preparing figures.

## A Proofs

**Proof of Proposition 1**. Using Equation (2), we can express *U*_*A,B*_(*x*) as

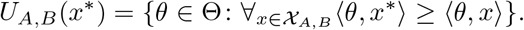

From the linearity of the inner product, ⟨*θ, x*^*^⟩ ≥⟨ *θ, x*⟩ is equivalent to ⟨*θ, x*^*^ −*x*⟩≥ 0. This allows us to write

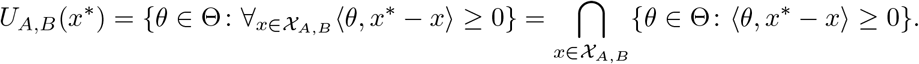

For any vector *y*, the set defined by the inequality ⟨*θ, y*⟩ ≥0 is a closed half-space, which is a convex set. Therefore, *U*_*A,B*_(*x*) is a closed convex set as an intersection of a finite number of such sets. Furthermore, since for any scalar *s* we have ⟨*sθ, y*⟩ = *s*⟨*θ, y*⟩, it follows that for *s >* 0 we have ⟨*sθ, y*⟩ ≥ 0 if and only if ⟨*θ, y*⟩ ≥ 0, which shows that *U*_*A,B*_(*x*) is a cone. The zero vector of parameters always belongs to *U*_*A,B*_(*x*), because ⟨0, *y*⟩ = 0 for any vector *y*. The proof for 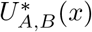 is similar; however, this set is open as an intersection of a finite number of open half-spaces. Furthermore, the zero vector never belongs to 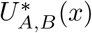,and the set may be empty. ◀

### Proof of Lemma 2

For the proof of Lipschitz continuity, consider two vectors of parameters *θ* and *θ*^*′*^. First, we note that

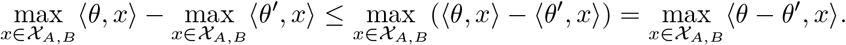

The inequality follows from the fact that

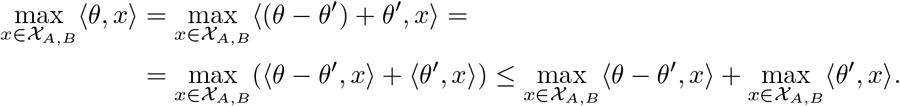

It follows that 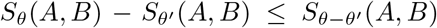.Next, using the Cauchy-Schwarz inequality ⟨*a, b*⟩ ≤ ||*a*|| · ||*b*||, we have

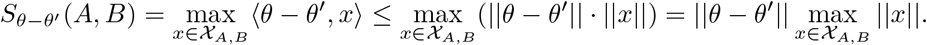

We therefore have 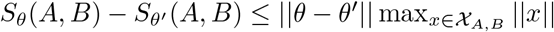. Swapping *θ* and *θ*^*′*^ (and noting that ||*θ* − *θ*^*′*^|| = ||*θ*^*′*^ = *θ*||), we obtain 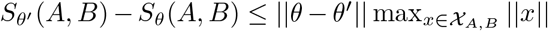, and thus we conclude that

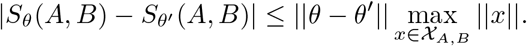

For the proof of convexity, consider *p*∈ [0, 1] and two vectors of parameters *θ* and *θ*^*′*^.

Then,

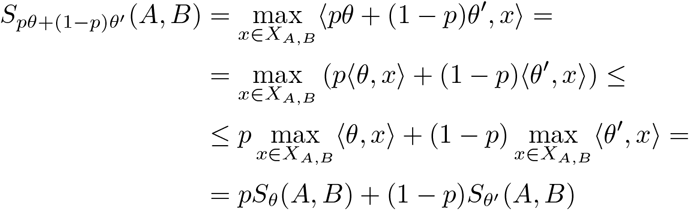

### Proof of Proposition 5

Since the functions *θ*→ *S*_*θ*_(*A*_*i*_, *B*_*i*_) is differentiable almost everywhere, so is *l*(*α, θ*;𝒟). Accordingly, let *θ* be such that every pair of sequences *A, B* has a unique optimal configuration *x*_*i*_, and let *U* = ⋂_*i*_ *U*^*^(*x*_*i*_) be the set of *θ* where all of these configurations are uniquely optimal. Then, the log-likelihood restricted to *U* can be expressed as

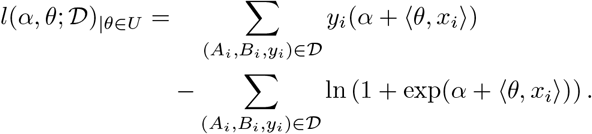

Now, 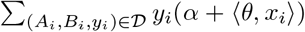 is linear with respect to *θ*, making it both convex and concave. Therefore, *l*(*α, θ*; 𝒟)_|*θ*∈*U*_ is concave with respect to *θ* as a sum of concave functions.

### Proof of Propostition 10

First, we observe that the functions ln *p*(*α, θ*; *A*_*i*_, *B*_*i*_), ln(1 −*p*(*α, θ*; *A*_*i*_, *B*_*i*_)) are locally Lipschitz continuous with respect to *θ*, so that their Clarke subdifferentials are well-defined. For a fixed intercept *α*, let *ς*(*x*; *α*): ℝ →ℝ be the logistic function of *x* defined as *ς*(*x*; *α*) = 1*/*(1 + exp( −*α* −*x*)). This function is globally Lipschitz continuous with respect to *x*, because for any *x* ∈ℝ we have *ς*^*′*^(*x*) = *ς*(*x*)(1− *ς*(*x*)) *<* 1. Next, since *p*(*α, θ*; *A*_*i*_, *B*_*i*_) = *ς*(*S*_*θ*_(*A*_*i*_, *B*_*i*_); *α*), and a composition of Lipschitz functions is Lipschitz, it follows that *p*(*α, θ*; *A*_*i*_, *B*_*i*_) is also globally Lipschitz continuous with respect to *θ*. Finally, ln(*x*) is locally (but not globally) Lipschitz continuous. The proof for ln(1 −*p*(*θ*)) follows the same lines. We remark that the function *l*(*α, θ*) is also locally Lipschitz with respect to *θ* as a sum of locally Lipschitz functions.

Now, the first equality in the Proposition follows directly from the chain rule in Lemma 9 applied to *f*(*θ*) = *S*_*θ*_(*A, B*) and *g*_*α*_(*x*) = − ln(1 + exp(−*α* − *x*)) for a fixed *α*. The second equality follows from the chain rule applied to *g*_*α*_(*x*) = − ln(1 + exp(*α* + *x*)). ◀

### Proof of Lemma 11

First, we note that from the definition of the Clarke subdifferential, we have 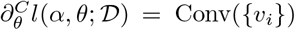 for a set of vectors *v*_*i*_, such that for each *v*_*i*_ there exists a sequence 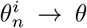 such that *l*(*α, θ*; *D*) is differentiable w.r.t. *θ* at each *θ*_*n*_ and 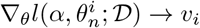.

Assume *θ*_*n*_ → *θ* is such that ℬ (*θ*_*n*_; 𝒟) = *{*(*x*_1_, …, *x*_|𝒟|_)*}* for all *n*. Then, for every pair of sequences *A*_*i*_, *B*_*i*_, the score *S*_*θ*_ (*A*_*i*_, *B*_*i*_) is differentiable w.r.t. *θ* at each *θ*_*n*_ (as the alignment configurations are unique). It follows that 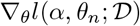 exists at all *θ*_*n*_. From Proposition 10, we get

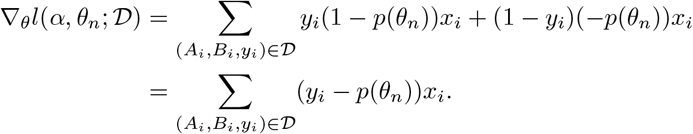

From the continuity of *p* it follows that 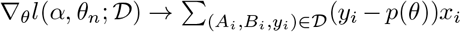, and thus 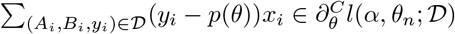.

We remark that the reason why the Lemma states an inclusion, rather than equality, is that a sum of non-differentiable functions can be differentiable (e.g. *f*(*x*) = |*x*| and *g*−(*x*) = |*x*| ). Therefore, even if the alignment scores are not differentiable at points of some sequence *θ*_*n*_→ *θ*, the log-likelihood potentially may be differentiable at these points and result in additional limits of gradients belonging to the Clarke subdifferential. Whether cases like that can happen in alignments remains an open question. ◀

### Proof of Theorem 14

First, we note that the log-likelihood function *l*(*α, θ*;𝒟 ) is a “tame” function in the sense of Davis, Drusvyatsky, Kakade and Lee and satisfies their Assumption D [10]. This follows from the fact that the function is composed of a finite number of algebraic operations, maxima over finite numbers of functions, and logarithmic and exponential functions, which makes it definable in the o-minimal structure ℝ_exp_ = (ℝ, +, ., exp, 0, 1) [61].

We will now show that the procedure satisfies Assumption C form [10]. The assumptions 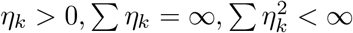correspond to Assumption C.1, and the assumption that *θ*_*k*_ is bounded corresponds to Assumption C.2. Now, let *α*_*k*_ = *ξ*_*k*_*/η*_*k*_, so that the update step can be written as *θ*_*k*+1_ = *θ*_*k*_ + *η*_*k*_(*g*_*k*_ + *α*_*k*_). Now, since 𝔼*ξ*_*k*_ = 0, we also have 𝔼*α*_*k*_ = 0. Furthermore, 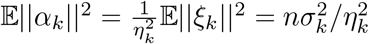. Therefore, *α*_k_ is a sequence of independent random variables with zero expected value and a bounded variance, which satisfies Assumption C.3. ◀

### Proof of Theorem 15

Let 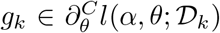.Let 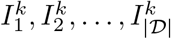 be a sequence of independent binary random variables such that 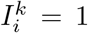 if the *i*-th pair was sampled in iteration *k*. By construction of 𝒟_*k*_, we have 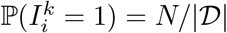.

Since the Theorem assumes that *g*_*k*_ is constructed using Proposition 12, conditioned on the sequence 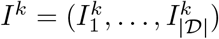 it can be expressed as

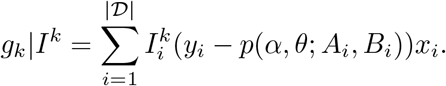

Let 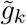 be a hypothetical subgradient obtained using Proposition 12 on the full dataset,

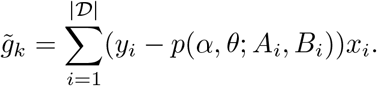

We now have

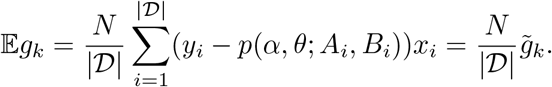

Let *α*_*k*_ = *ξ*_*k*_*/η*_*k*_ and let 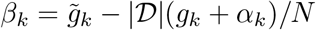 . We can now write the *θ* update step as

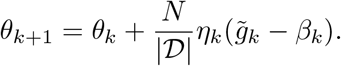

The Theorem now follows from Theorem 14 applied to the above sequence, treating *Nη*_*k*_*/*|𝒟| as the sequence of step lengths and *β*_*k*_ as the sequence of random errors.

